# Association Between The Posterior Part Of The Circle Of Willis And Vertebral Artery Hypoplasia

**DOI:** 10.1101/557397

**Authors:** Virginija Gaigalaite, Jurate Dementaviciene, Augenijus Vilimas, Danute Kalibatiene

## Abstract

**Background:** It is not clear whether the configuration of the posterior part of the circle of Willis (CW) depends on the proximal part of the vertebrobasilar system. Our aim is to evaluate the posterior part of CW in association with different size of vertebral arteries (VA) in subjects free from stroke and TIA.

**Materials and methods:** The present study was based on a sample of 923 subjects free from stroke and TIA who were examined from 2013 through 2018. All the participants underwent MRA examination. The duplex ultrasonographic examination of the extracranial arteries (vertebral and carotid) was performed. VA was defined as hypoplastic (VAH) when VA diameter in the entire course was less than 2.5 mm. We classified the posterior communicating arteries (PCoA) as presence PCoA, absence/hypoplastic PCoA and fetal CW (FCW). The comparison of the posterior part of CW was made in subjects with normal VA and VAH of a different degree (communicating with basilar artery (VAH-BA) and not communicating with the basilar artery and terminating in PICA, neck or aplasia (VAH-PICA)).

**Results:** FCW was found in 15.9% of subjects, bilaterally – in 2.3 %. The coexisting VAH was more common in subjects with FCW rather than in those with adult CW (respectively, 28.6% and 13.4%, p<0.001). Aplasia of A1 of the anterior cerebral artery, i.e. blood flow redistribution in the anterior part of anterior circulation in the majority of cases (in 6 of 7 cases) was found ipsilaterally to FCW. FCW was recorded in 50% of the subjects with VA - PICA in comparison with 13.5% of those with normal VA and 22.8% with VAH - BA, p<0.005. On the contrary, absence/hypoplasia of both PCoA was mostly found in the group with normal VA in comparison with VAH-BA and VAH-PICA (accordingly, 50.7%, 38.6% and 12.5%, p<0.01).

**Conclusion:** Individuals with VAH have a different pattern of the posterior part of CW in comparison with those with normal VA. With the increasing degree of VAH, the proportion of FCW increases, while the proportion of absence/hypoplastic of both PCoA decreases.

## Introduction

The Circle of Willis (CW) is a major intracranial collateral circulation that has an important role in ischemic events. The most common configuration of the posterior part of CW is described as ‘adult configuration’. In these cases, the posterior cerebral artery (PCA) is a terminal branch of the vertebrobasilar system. The diameter of the precommunicating part (P1) of PCA is larger than the diameter of the posterior communicating artery (PCo A) connecting the vertebrobasilar and carotid systems. The presence of PCoA enables to redistribute the blood flow in both directions through PCoA in cases of diminished blood supply in the internal carotid artery (ICA) or vice versa in the vertebrobasilar system. In the minority of cases the configuration of the posterior part of CW is the so called fetal-type of the posterior circle of Willis (FCW). FCW is a morphological variant of the cerebrovascular anatomy in which PCA arises directly from the terminal ICA, with or without an intact P1 segment connecting PCA to the basilar artery. In this variant, the larger brain area is dependent on ICA and could be more prone to develop large ischemic strokes in cases of carotid artery stenosis or occlusion. As described by many authors [1], in these cases the collateral circulation between the anterior and posterior circulation through secondary collaterals, i.e. leptomeningeal vessels cannot develop since both, the middle cerebral artery and PCA are connected to the same internal carotid system.

Insufficient attention has been given to the FCW coexistence with other vascular congenital variants and its influence on both cerebral circulation and neurological symptomatic. It was described [2] that the coexistence of FCW, basilar artery (BA) hypoplasia and vertebral artery hypoplasia (VAH) was more common in patients with cerebral ischemia, i.e. this arterial variant may increase TIA/stroke risk. According to [3], individuals with FCW have an 18% reduction in BA diameter. It is not clear if the configuration of the posterior part of CW depends on the proximal part of the vertebrobasilar system, more exactly, on the vertebral artery (VA) diameter and in cases of a small diameter vertebrobasilar system, which configuration of posterior collateral circulation is more beneficial.

Our aim is to evaluate the posterior part of CW in association with different sizes of VA (normal diameter, VAH of different degree (communicating with the basilar artery (VAH-BA) and not communicating with the basilar artery and terminating in PICA, neck or aplasia (VAH-PICA)) in subjects free from stroke and TIA.

## Material and methods

The present study was based on a sample of 923 subjects without cerebrovascular disease (TIA or stroke) history before and at the time of the study enrollment. All of them were examined by magnetic resonance imaging (MRI) and magnetic resonance angiography (MRA) in the Republican Vilnius University Hospital from 2013 through 2018. The inclusion criteria were as follows: (1) no history of transient ischemic attack, ischemic or hemorrhagic stroke; (2) no disabling neurological deficits on examination; (3) extracranial or intracranial vessels without significant stenosis (>50%) or occlusion; (4) the study excluded the patients who did not undergo MRI or MRA investigation, their intracranial vessels were not visualised or they refused to participate in the study.

### Imaging studies

All the participants underwent MRA examination using 1.5 Tesla MRI (GE Optima MR450w 1.5T MRI System) for the brain and CW evaluation.

The following sequences were obtained: 3D T1 weighted, T2 FLAIR, T2 weighted, diffusion weighted imaging (DWI, b-0, b-1000), SWAN (Susceptibility weighted Angiography), 3D Time of Flight MR angiography (3D-TOF-MRA). CW anatomy of each individual was evaluated using both 3DTOF MRA MIP reconstructions and source images.

The duplex ultrasonographic examination of extracranial arteries (vertebral and carotid) was performed by using the 7.5 MHz linear array transducer of Aloka Prosound F 75 ultrasound system. The diameter of VA in our previous study was measured similarly [4].

### Image analysis

MRA were reviewed by two independent neuroradiologists. If they had disagreements regarding the configuration of the circle of Willis, they discussed it until a consensus was reached.

The classification of CW and VA was carried out as follows:

1. When interpreting MRA, the presence or absence of PCcoA and P1 segment of PCA was assessed. P1 segment and the posterior communicating artery (PCoA) were scored as normal (diameter ≥0.8 mm), hypoplastic (diameter <0.8 mm in MRA), absent or non-visualised. The threshold of 0.8 mm in MRA was chosen in order to be consistent with other studies reported in literature [5].
2. The posterior part of the circle of Willis was defined as complete in cases of presence of both PCoA and P1 segment of PCA with diameter ≥0.8 mm. All other variants were defined as the incomplete posterior part of CW.
3. We defined the circle of Willis as fetal if PCA arises from the internal carotid artery, independent on the presence or absence of the atretic P1 segment. All other individuals were named as having” adult” configuration of CW. The subjects with adult and rarely found transitional CW configuration were included in this group. In cases of adult configuration, P1 segment of PCA had a diameter larger than PCoA while in transitional configuration, P1 segment and PCoA have close diameters.
4. The posterior part of CW was documented as presence of PCoA, absence/hypoplastic PCoA and FCW. The subjects with hypoplastic PCoA were included in the same group as those with absence of PComA since both groups have minimal or no possibilities to compensate the reduced posterior circulation from carotid arteries through PCoA in comparison with the individuals with presence of normal PCoA.
5. The absence of A1 segment of the anterior cerebral artery (ACA) was documented. In these cases both A2 segments are supplied by the existing A1 from the contralateral ACA.
6. VAH was established according to MRA (V4 segment) and duplex scanning (V1-V3 segment). We defined VA as hypoplastic when VA diameter in the entire course was less than 2.5 mm. We also studied the group named “VAH-PICA” with VA aplasia or hypoplastic VA not communicating with BA and terminating in PICA or the neck, i.e. subjects with the possibility of the greatest reduced blood flow through VA.

We present a pattern of the posterior part of CW found in our individuals. We have estimated what concomitant vascular variants of the vertebrobasilar system are more common in FCW in comparison with the adult CW. Also, we have estimated if FCW influence blood flow redistribution in the anterior part of anterior circulation, i.e. if both A2 segments are supplied by the existing A1 from the contralateral or ipsilateral sides of ACA.

We assessed if the posterior part of CW differs in subjects with normal VA, VAH communicating with BA and VAH not communicating with BA or VA aplasia.

### Statistics

The *Chi* square independence (χ2) test was applied in carrying out the comparison between the categorical variables, while Fisher’s exact test was used in the case of a small sample size. Continuous variables meeting the assumptions of normality were analysed using t-tests for independent groups. The chosen significance level was α=0.05.

#### Ethics

This study was approved by the Ethics Committee for the Vilnius region (No. 158200-15-767-281).

## Results

### Posterior part of CW

The characteristics of the posterior part of CW are presented in Table 1. FCW was found in 15.9 % of subjects free of stroke and TIA. Side-related differences in the posterior part of CW observed in both types of CW did not reach a statistically significant difference. In 47.9% of individuals both PCoA were absent or hypoplastic. 2.3 % of the subjects had both-sided FCW.

**Table 1.** The characteristics of the posterior part of the circle of Willis (n=923)

| Type of posterior part of CW | N | Proportion (%) |
| --- | --- | --- |
| Adult type | 776 | 84.1 |
| • Absence of both PCoA | 442 | 47.9 |
| • Absence of one PCoA | 191 | 20.7 |
| ○ Absence of left PCoA | 112 | 12.1 |
| ○ Absence of right PCoA | 79 | 8.6 |
| • Both PCoA | 143 | 15.5 |
| Fetal type | 147 | 15.9 |
| Side of fetal circle of Willis |  |  |
| • Left-sided | 58 | 6.3 |
| • Right-sided | 68 | 7.4 |
| • Both-sided | 21 | 2.3 |
| Contralateral PCoA |  |  |
| • Absence/hypoplasia | 81 | 8.8 |
| • Normal | 45 | 4.9 |
| • Fetal circle of Willis (both-sided) | 21 | 2.3 |

*Demographic characteristics and coexisting arterial variants in adult CW and FCW* (Table 2). The proportion of men and women did not differ in both configurations of CW. The coexisting VAH was more common in subjects with FCW than in subjects with adult CW (correspondingly, 28.6% and 13.4%, p<0.001). Aplasia of A1was rare in both groups, although aplasia of A1 was more common in the group with FCW compared to those with adult CW. Moreover, in the majority of the subjects with FCW (in 6 of 7 cases), A1 aplasia was found ipsilaterally to FCW, the carotid artery supplies blood to PCA and MCA, while ACA is receiving blood from the contralateral carotid artery.

**Table 2.** The comparison of coexisting characteristics in FCW and adult CW

|  | Fetal circle of Willis<br>N=147 | Adult circle of Willis<br>N=776 | p-value |
| --- | --- | --- | --- |
| *Demographics:* |  |  |  |
| Men | 79 (53.7%) | 448 (57.7%) | NS* |
| Age | 46.4±1.4 | 48.1±0.4 | NS |
| *Vertebrobasilar system* |  |  |  |
| VAH | 42 (28.6%) | 104 ( 13.4%) | <0.001 |
| VAH terminating<br>in PICA/aplasia | 16 (10.9%) | 16 ( 2.1%) | <0.001 |
| BA hypoplasia<br>/aplasia | 2 (1.4%) | 5 (0.64%) | NS |
| Fenestration of BA | 1 (0.7%) | 3 (0.4%) | NS |
| *Anterior part:* Aplasia<br>of A1 (ACA artery)<br>segment and both A2<br>segments are supplied<br>from one side by the<br>existing A1 | 7 (4.8%)<br>Ipsilateral side: 6 (4.1%)<br>Contralateral side: 1<br>(0.7 %) | 13 (1.7%) | 0.018 |

\*NS-p>0.05

### Association between VAH and the pattern of the posterior part of CW

The association between VAH and the variants of the posterior part of CW is presented in Table 3. The pattern of the posterior part of CW in subjects with VAH differs from those with normal VA. FCW was more frequent in individuals with VAH than in those with normal VA (accordingly, 28.8 % vs. 13.5%, p<0.001), while the absence/hypoplasia of both PCoA was more common in subjects with normal VA in comparison to those with VAH (accordingly, 50.7% and 32.9%, p<0.001).

**Table 3.**
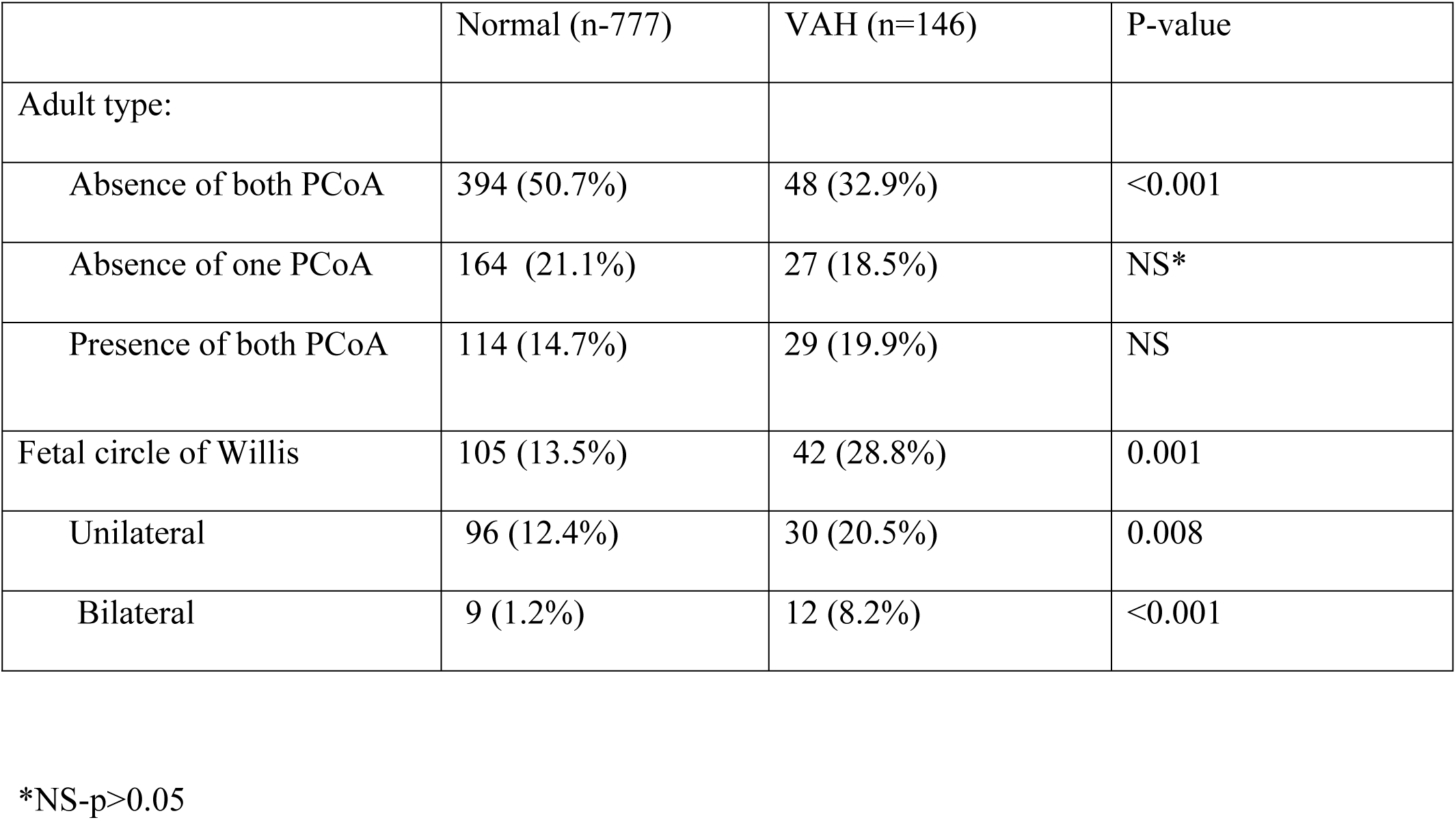
The comparison of the posterior part of CW in patients with VAH and normal VA.

|  | Normal (n=777) | VAH (n=146) | P-value |
| --- | --- | --- | --- |
| Adult type: |  |  |  |
| Absence of both PCoA | 394 (50.7%) | 48 (32.9%) | $<0.001$ |
| Absence of one PCoA | 164 (21.1%) | 27 (18.5%) | NS* |
| Presence of both PCoA | 114 (14.7%) | 29 (19.9%) | NS |
| Fetal circle of Willis | 105 (13.5%) | 42 (28.8%) | 0.001 |
| Unilateral | 96 (12.4%) | 30 (20.5%) | 0.008 |
| Bilateral | 9 (1.2%) | 12 (8.2%) | $<0.001$ |

\*NS- $p>0.05$

### Difference of the posterior part of CW in subjects with VAH not communicating with the basilar artery and those with VAH communicating with the basilar artery

The above mentioned regularity was even more striking in the least developed hypoplastic VA that do not communicate with the basilar artery (Table 4). Half of the patients with VAH - PICA had FCW compared to 13.5% of those with normal VA diameter and to 22.8% of individuals with VAH that communicates with the basilar artery, p<0.005. Moreover, the proportion of bilateral FCW was largest in the subjects with VAH-PICA. On the contrary, the absence of both PCoA was most frequent in the group with normal VA and rare in VAH-PICA group (accordingly, 50.7% and 12.5%, p<0.001).

**Table 4.** Comparison of the posterior part of CW in patients with a different degree of VAH.

|  | Normal (n=777) | VAH-BA (n=114) | VAH-PICA (n= 32) | p-value |
| --- | --- | --- | --- | --- |
| Adult type: |  |  |  |  |
| Absence of both PCoA | 394 (50.7%) | 44 (38.6%) | 4 (12.5%) | 0.001 |
| Absence of one PCoA | 164 (21.1%) | 22 (19.3%) | 5 (15.6%) | NS |
| Presence of both PCoA | 114 (14.7%) | 22 (19.3%) | 7 (21.9%) | NS |
| Fetal circle of Willis | 105 (13.5%) | 26 (22.8%) | 16 (50%) | 0.001 |
| Unilateral | 96 (12.4%) | 20 (17.5%) | 10 (31.3%) | 0.004 |
| Bilateral | 9 (1.2%) | 6 (5.3%) | 6 (18.8%) | 0.001 |

### The sides of VAH and FCW

VAH and FCW were more frequently observed on the same side: VAH was observed ipsilaterally to FCW in 76% of cases, VAH - PICA - ipsilaterally to FCW in 82.4% of cases.

## Discussion

FCW was found in 15.9% of individuals, bilaterally - in 2.3% of cases. According to other authors, the proportion of FCW ranges from 11 to 32% [1], [6], [7], [8], [9]. VAH was observed in 12.5 % of subjects. According to the data presented by other authors, depending on the VAH definition, the method of examination and the category of population, this proportion ranges from 1.9% to 25% [10]. Therefore, the population under our investigation was a typical population.

FCW was more frequently observed in subjects with VAH, i.e. with an insufficiently developed proximal part of the vertebrobasilar system, compared to those with normal VA diameter. Among individuals with a very small VA terminating in PICA/neck/aplasia, compared to those subjects whose VA is wider and forms the basilar artery, the proportion of FCW was larger. Moreover, FCW was more common in ipsilateral to VAH side rather than contralateral.

According to [3], BA diameter is inversely associated with FCW. Otherwise, FCW was more common in individuals with an insufficiently developed distal part of the vertebrobasilar system which can lead to inadequate posterior circulation, rather than in those with the normal basilar artery. The influence of an insufficiently developed proximal part of the vertebrobasilar system, including VAH or aplasia on the posterior circulation insufficiency and as a consequence the demand to compensate the possible inadequate blood supply to the brain is under discussion. Although, many authors estimate VAH as an independent predictor of stroke or TIA [10]. The hypothesis that VAH can lead to the posterior circulation insufficiency is also supported by our results that with the decreasing VA diameter, the risk of stroke/TIA increases [4]. Moreover, VAH can lead to a relative regional hypoperfusion in the PICA territory [11]. As described in a study [1], during the embryological development the anterior circulation supplies the occipital region, the brain stem and the cerebellum via multiple anastomoses because the posterior circulation is not yet well developed. After the development of VA and sufficient posterior circulation, these anastomoses regress. FCW as a result of failed regression that may be associated with insufficient blood supply via the insufficiently developed vertebrobasilar system, including hypoplastic. In these cases the carotid artery may particularly recall the role of the vertebrobasilar system by supplying blood to the posterior fossa as in the embryological development. The greater proportion of FCW in subjects with more severe VAH whose blood flow through VA is reduced to a greater degree supports the hypothesis that with the decreasing blood supply from VA to the brain, the possible inadequate perfusion in posterior circulation is more frequently compensated through FCW from anterior circulation, ICA. In case of small diameter VA, compared to normal diameter VA, FCW may provide better blood supply to the brain and prevent from cerebral ischemia.

In summary, the pattern of the posterior part of CW in stroke/TIA-free subjects with VAH and normal VA was different. The proportion of absence/hypoplasia of both PCoA, i.e. the absence /hypoplasia of primary collaterals was larger in subjects with normal proximal circulation, i.e. normal VA diameter compared to those with insufficiently developed proximal part of the vertebrobasilar system, VAH. And vice versa, the proportion of FCW was larger in those with VAH compared to those with normal VA diameter. The proportion of subjects with a complete posterior part was larger in those with VAH although the difference did not reach a statistically significant difference. These results support the hypothesis that in cases of small vertebral arteries the collateral circulation through PCoA or FCW may be important for prevention of stroke/TIA in the posterior circulation. Future investigations are needed in order to assess whether in cases of VAH the configuration of the posterior part of CW can prevent or increase the stroke/TIA risk. The study [1] revealed that the coexistence of the basilar artery hypoplasia, VAH and the fetal CW were more common in stroke patients. However, in the above mentioned study, the role of FCW is not clear. Is FCW an independent stroke predictor, or is FCW not able to compensate the reduced blood flow in cases of coexistence of small proximal and distal parts of the vertebrobasilar system? Future investigations are needed on the associations between a small vertebrobasilar system, CW configuration and neurological symptoms such as vertigo. Patients with stroke/TIA were excluded from our study, however, suggestions can be made for further studies to compare how CW differs in vertigo patients and healthy subjects. In cases of FCW, the territory supplied with blood by the carotid artery increases up to three arteries (ACA, middle cerebral artery (MCA) and PCA). A1 aplasia in most cases is found ipsilateral to FCW and may be associated with the need to redistribute the blood flow in the anterior circulation and to reduce the territory of blood supply from the carotid artery from three arteries territory (ACA, MCA, PCA) to two arteries territory (MCA, PCA, while both ACA are supplying from the contralateral carotid artery.

## Conclusions

Individuals with VAH have a different pattern of the posterior part of CW in comparison with those with normal VA diameter. With the increasing degree of VAH, the proportion of FCW increases while the proportion of absence/hypoplasia of both PCoA decreases.

## Acknowledgements

We thank Leonora Norviliene for providing assistance in the revision and translation of the manuscript.

